# FILER: large-scale, harmonized FunctIonaL gEnomics Repository

**DOI:** 10.1101/2021.01.22.427681

**Authors:** Pavel P. Kuksa, Prabhakaran Gangadharan, Zivadin Katanic, Lauren Kleidermacher, Alexandre Amlie-Wolf, Chien-Yueh Lee, Liming Qu, Emily Greenfest-Allen, Otto Valladares, Yuk Yee Leung, Li-San Wang

## Abstract

**Motivation:** Querying massive collections of functional genomic and annotation data, linking and summarizing the query results across data sources and data types are important steps in high-throughput genomic and genetic analytical workflows. However, accomplishing these steps is difficult because of the heterogeneity and breadth of data sources, experimental assays, biological conditions (e.g., tissues, cell types), data types, and file formats.

**Results:** FunctIonaL gEnomics Repository (FILER) is a large-scale, harmonized functional genomics data catalog uniquely providing: 1) streamlined access to >50,000 harmonized, annotated functional genomic and annotation datasets across >20 integrated data sources, >1,100 biological conditions/tissues/cell types, and >20 experimental assays; 2) a scalable, indexing-based genomic querying interface; 3) ability for users to analyze and annotate their own experimental data against reference datasets. This rich resource spans >17 Billion genomic records for both GRCh37/hg19 and GRCh38/hg38 genome builds. FILER scales well with the experimental (query) data size and the number of reference datasets and data sources. When evaluated on large-scale analysis tasks, FILER demonstrated great efficiency as the observed running time for querying 1000x more genomic intervals (10^6^ vs. 10^3^) against all 7×10^9^ hg19 FILER records increased sub-linearly by only a factor of 15x. Together, these features facilitate reproducible research and streamline querying, integrating, and utilizing large-scale functional genomics and annotation data.

**Availability and implementation:** FILER can be 1) freely accessed at https://lisanwanglab.org/FILER, 2) deployed on cloud or local servers (https://bitbucket.org/wanglab-upenn/FILER), and 3) integrated with other pipelines using provided scripts.

## 1. Introduction

Functional genomic data and annotations are commonly used to provide the necessary functional evidence in various systems biology, genetic and genomic analyses, such as the analysis of the non-coding genome-wide association study (GWAS) signals or the analysis of the experimentally-derived genomic regions (Kuksa *et al.*, 2020; Amlie-Wolf *et al.*, 2018; Watanabe *et al.*, 2017; Nagraj *et al.*, 2018; Rouillard *et al.*, 2016; Dozmorov *et al.*, 2016). Such functional genomic annotation includes different types of data such as tissue-specific regulatory elements (enhancers) (Andersson *et al.*, 2014), transcription factor (TF) binding activity (Dunham *et al.*, 2012; Davis *et al.*, 2018; Heinz *et al.*, 2010), chromatin states (Kundaje *et al.*, 2015; Song and Crawford, 2010), genetic regulation (expression quantitative trait loci (eQTL), splicing QTLs (sQTL)) information (Aguet *et al.*, 2017, 2020) and chromatin conformation data (Lieberman-Aiden *et al.*, 2009). These data originate from a variety of sample sources including primary tissues, primary cells, immortalized cell lines, in vitro differentiated cells and others.

The primary functional genomic experimental data is made available by many major consortia such as ENCODE (Davis *et al.*, 2018; Dunham *et al.*, 2012), GTEx (Aguet *et al.*, 2020, 2017), FANTOM5 (Andersson *et al.*, 2014), and NIH Roadmap Epigenomics (Kundaje *et al.*, 2015). So far, these four projects generated datasets spanning over >50,000 experiments across >1,000 tissues, cell types, biological conditions, with each dataset containing millions to billions of annotation or assay readout records across the genome. In order to pair these functional annotations with high-throughput analytical workflows, e.g., for processing current population-level studies such as UK Biobank (Bycroft *et al.*, 2018) (500,000 individuals with >2,500 phenotypes), we need a scalable, unified, high-throughput and robust access to these massive, heterogeneous genomic data collections.

To address these issues, we developed FunctIonaL gEnomics Repository (FILER), a large-scale, curated, integrated catalog of harmonized functional genomic and annotation data coupled with a scalable genomic search and querying interface to these data. The latest FILER release provides seamless integration of >58,000 harmonized genomic datasets (*data tracks*) across diverse (>20) primary data sources organized into >140 data collections, wide biological context (>1,100 cell types) and various genomic and biological features (>30 experimental assays and data types). FILER provides a unified access to this rich functional and annotation data resource spanning >17 Billion records across genome with >2,700x total genomic coverage for both GRCh37/hg19 and GRCh38/hg38. FILER can be used to perform flexible querying, staging and consolidation of these data for analyses. FILER is accessible via a web server (https://lisanwanglab.org/FILER) or can be deployed on users’ own cloud computing instances, local servers, high-performance computing clusters (https://bitbucket.org/wanglab-upenn/FILER) and integrated with genomic and genetic analysis workflows.

## 2. Methods

### 2.1. FILER structure and data organization

FILER has been implemented using an easily updatable, extensible, and modular architecture. **Figure 1** gives an overview of the FILER architecture and data organization. FILER contents are derived from the integration of many data and annotation resources as well as curation and processing of publicly available functional genomics datasets (**Figure 1A**). **Supplementary Table S2** (’Data sources’), and **Supplementary Table S3** (’Data collections’) show a detailed list of data sources and data collections available in FILER. First, primary annotation and functional genomics datasets (referred to as *data tracks*) from existing data sources are collected and compiled into a unified catalog, as shown in the diagram in **Figure 1A**. Second, individual genomic datasets are curated, processed, and imported into FILER (**Figure 1A** diagram) using the FILER data harmonization and annotation pipeline (see **Section 2.2** for details). Data-source-specific metadata schemas were matched across data sources with the FILER schema (**Supplementary Table S5** provides details of the schema matching) to generate standardized, consistent meta-data descriptions for each of the FILER data *tracks* (sets of genomic/annotation records).

**Figure 1.**
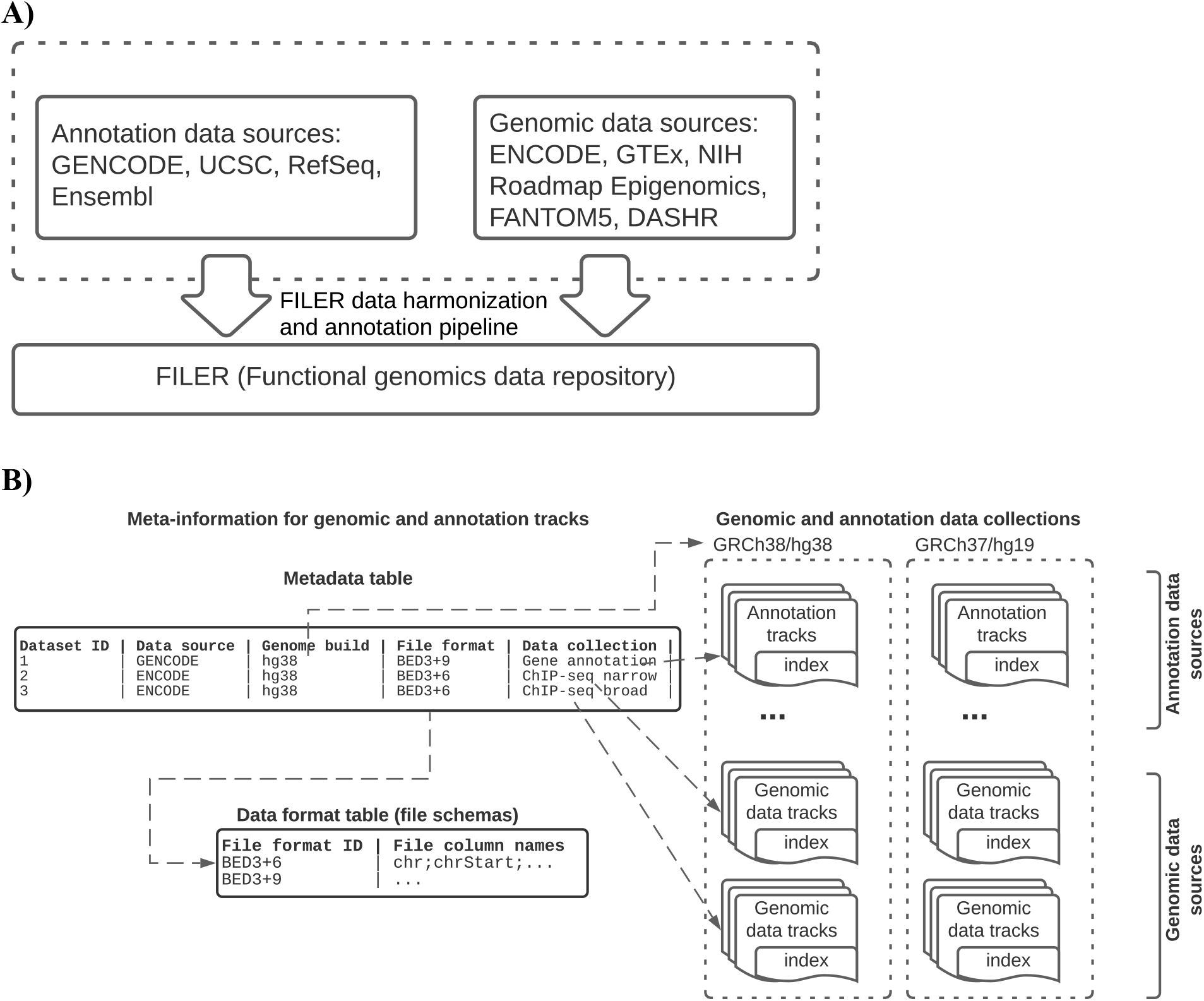
(**A**) Overview of the FILER architecture. Annotation resources and genomic data sources are collected and compiled into a unified data catalog. All datasets are curated, systematically processed, and imported into FILER using the data harmonization and annotation pipeline (see **Section 2.2**, **Figure 2** for details). (**B**) Organization of FILER. Metadata and file schemas tables (shown on the left-hand side) contain detailed, standardized meta-information for individual datasets and descriptions of the data file schemas.

All datasets in FILER are organized into data collections (shown on the right-hand side) by data source, experimental assay, data type, file format, and genome build (see **Section 2.1** for details on FILER data organization). For each FILER dataset, the metadata table contains a pointer to the data collection containing the dataset (pointers are shown with dashed arrow lines in the figure). Data collections are indexed to enable efficient search and retrieval.

Each data source available in FILER (e.g., ENCODE, Roadmap, GTEx) is organized into one or more *data collections* according to the experimental assay, data type/file format, and the genome build. This ensures that each data collection only contains tracks sharing the same file format, the same genome assembly, and the same experimental protocol, thus allowing all such tracks to be indexed together and, importantly, allowing query (e.g., overlap) results to be combined across tracks. Some data collection examples in FILER include ENCODE ChIP-seq called peaks in the narrow peak format, ENCODE DNase-seq peaks, or called small RNA loci from the DASHR database (Kuksa *et al.*, 2019; Leung *et al.*, 2016; Kuksa *et al.*, 2018) of small non-coding RNAs across tissues/cell types. **Section 2.2** describes FILER data collections and organization in more details. **Supplementary Tables S2, S3** provide details on data sources and data collections available in FILER.

Internally, FILER stores information in several tables:

1. meta-information table (**Figure 1B; Supplementary Table S4**) containing the standardized information for each data track,
2. file schema table (**Figure 1B**), and
3. external, indexed data collections holding the actual genomic data (right-hand side of **Figure 1B**) referenced in the meta-information table.

We describe each of these tables in the next sections below.

#### 2.1.1. FILER meta-information table

The FILER metadata table stores detailed information for each data track including the data source (e.g., ENCODE, Roadmap, FANTOM5), experimental assay (e.g., ChIP-seq, DNase-seq, ATAC-seq), type of genomic records (e.g., called peaks, transcription start sites, gene models), biological source (e.g., cell type, tissue, cell line), data provenance (e.g., the data source version (if applicable), the date data was accessed/downloaded, download URL). Both the original (as obtained from the data source) data files and the processed files are stored in FILER in separate folder locations. The metadata table store the references to the original file including the download URL and the local file location. Each data track is uniquely identified by a FILER identifier assigned when the data is added to FILER (see, e.g., **Figure 1B** and **Supplementary Table S8**).

Additionally, the metadata table stores a number of derived and computed properties used for data integration and organization purposes, including tissue category and data category. Tissue category is used to define terms corresponding to individual cell types/tissues into a broader, standardized tissue/cell type category (see **Supplementary Methods,**section on tissue categorization). The data category is designed to provide a biologically meaningful description of each track. The data category for each track in each of the data sources is systematically generated based on a combination of the data attributes, including the experimental assay (e.g., ChIP-seq, DNase-seq), experimental target (antibody, if applicable; e.g., CTCF protein, or H3K27ac histone mark), and the type of genomic records (e.g., narrow peaks, transcription start sites) (see **Supplementary Methods; Section 2.2)**on data track classification/categorization for details).

#### 2.1.2. FILER file schema information table

The data tracks in FILER are stored in formats specific to particular data sources and/or experimental assays. The file schema table stores the schemas (see Figure 1B bottom) for each type of data tracks. These file schemas can be used to extract additional information from the genomic records contained in the track files. Each record in the file schema table defines the number of standard BED fields, the number of extra fields, as well as the names of all the fields in the genomic record (see **Figure 1B** (bottom left) for an example; for more detailed information on the FILER database schema table please refer to **Supplementary Table S6)**.

#### 2.1.3. FILER indexed data collections

The external indexed data collections (see **Figure 1B**, right panel) are referenced in the FILER metadata table using pointers to the file directories holding the genomic/annotation data for each of the FILER data collections. Data indexes are created for each data collection and are used to accelerate access/search within data collections. The data indexes for each of the data collections are located within the data collection file directories in the standardized location (the `‘giggle_index’` sub-folders). **Supplementary Table S3** provides details on the indexed data collections available in FILER. Data collections in FILER thus are stored in separate folders with each folder holding both the actual genomic datasets the corresponding genomic data index.

### 2.2. FILER data harmonization and annotation pipeline

The main steps in the FILER data harmonization and annotation pipeline (**Figure 2**) include 1) data annotation (metadata extraction and generation), 2) data pre-processing and normalization, 3) data classification and organization into data collections, and 4) genomic interval-based data collection indexing (Layer *et al.*, 2018).

**Figure 2.**
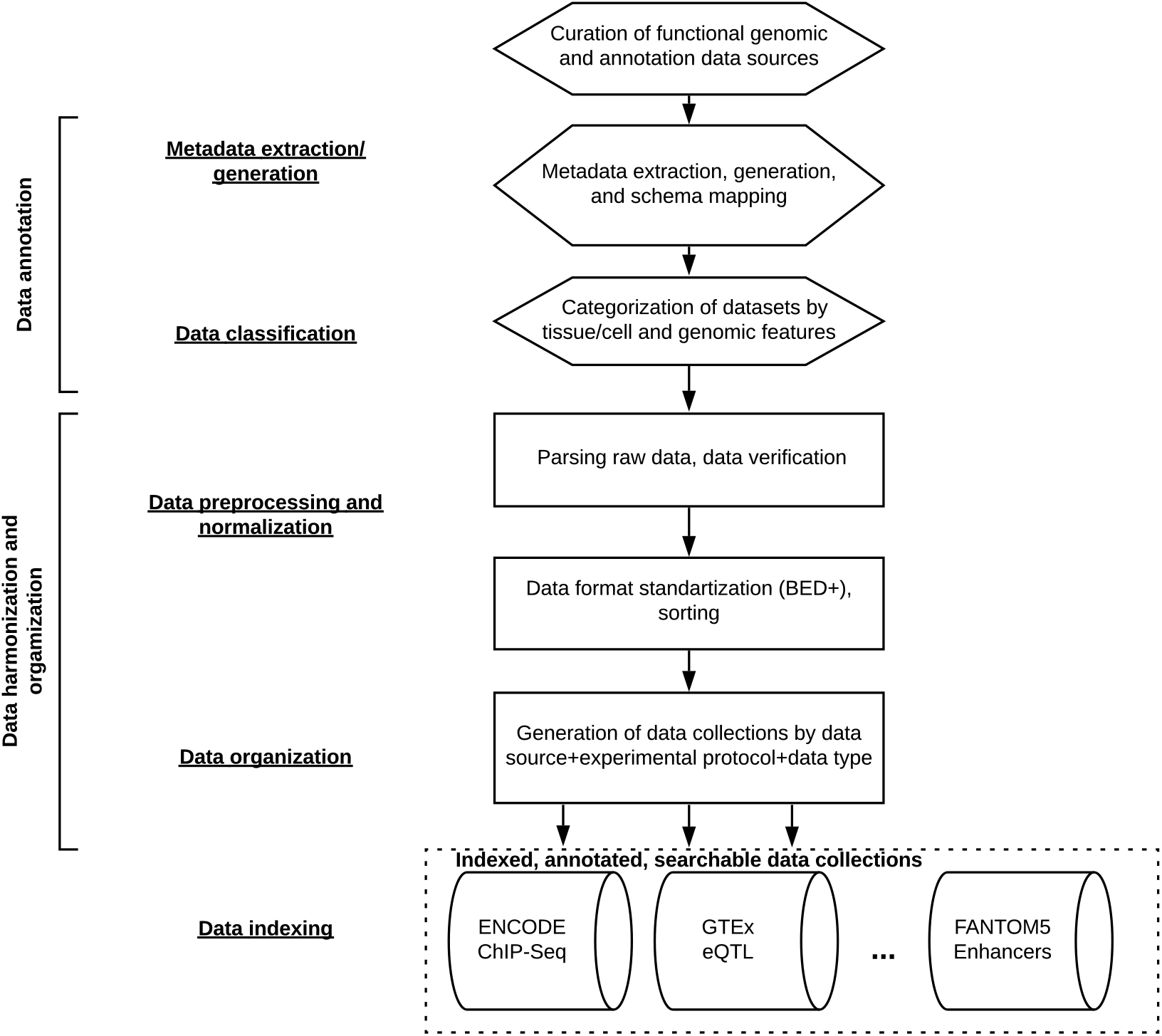
FILER data annotation and harmonization pipeline. All functional genomic datasets from the curated data sources are annotated with a standardized, consistent meta-information to allow for integration across diverse data sources (see **Sections 2.1, 2.2**). All individual genomic datasets are systematically pre-processed into standard, uniform BED-like file formats, and grouped into data collections based on the data source, experimental data type, data format (see **Section 2.1**). Each of the resulting data collections is indexed to allow efficient access using genomic-interval based indexing and query engine (see **Section 2.1** for details).

#### 2.2.1. Data annotation

All genomic tracks were annotated with the standard set of attributes (meta-information) including data type (ChIP-seq peaks, transcription factor binding sites (TFBS), gene models, etc), assay type (ChIP-seq, DNAse-seq, ATAC-seq, etc), cell/tissue type, data source, data source version, and other relevant meta-information (see **Supplementary Table S4** describing FILER metadata schema). To extract/infer the necessary meta-information for each data track, the information provided in the original data source was parsed and matched to the standardized schema employed by FILER (see **Supplementary Table S5** for details).

#### 2.2.2. Data pre-processing

Uniform pre-processing (format standardization, file schema inference and normalization, genomic coordinated-based sorting, compression, indexing) is performed to convert each original data track (e.g, in GFF (GFF3), BigBed (Kent *et al.*, 2010) formats) into a standard, BED-like format. **Supplementary Table S6** describes the file schemas for all data file formats integrated into FILER. Additionally, conversion of the genomic coordinates (liftover (Kuhn *et al.*, 2013)) from GRCh37/hg37 to GRCh38/hg38 was performed for datasets available exclusively in the earlier GRCh37/hg19 genomic build. Currently, FILER includes genomic tracks for GRCh37/hg19 (hg19) and GRC38hg38 (hg38, hg38-lifted) genomic builds.

#### 2.2.3. Data classification and organization into data collections

To further organize data and enable efficient search and retrieval of the functional genomics and annotation tracks, datasets from the same data source, same assay, same type of genomic records, same data format were physically grouped into data collections by data source, assay type, data type, and genome build (see **Supplementary Table S3** for details of data collections available in FILER). Each data collection is stored in a separate directory and contains files in the same format (e.g., BED3, BED6, etc) and is indexed using genomic interval-based indexing (Layer *et al.*, 2018) (see **Section 2.2.4 ‘Genome interval-based data indexing’** for details on data indexing in FILER). This genomic interval-based indexing schema created for each genomic collection facilitates cross-data collection search (e.g., genomic interval-based query) in FILER.

Additionally, all genomic and annotation data tracks included in FILER were categorized based on the biological source, tissue/cell type into broader tissue/cell type categories to further enable cross-data source integration of the data tracks (see **Supplementary Methods**, Tissue categorization section for details of tissue/cell type-based categorization employed by FILER). All FILER data tracks were annotated with a major tissue and organ systems category they belong to. FILER tracks were also systematically classified into biologically meaningful data categories (**Section 2.1.1**) to allow the users to more easily access datasets of interest.

#### 2.2.4. Genomic-interval based data indexing

Each data collection in FILER (**Supplementary Table S3; Section 2.2.3**) is indexed using a customized genomic-interval based indexing (Layer *et al.*, 2018). Indexing creates B+ tree-like structures over genomic intervals contained in the data collection. The leaf nodes contain pointers to the data files and file offsets for individual genomic records (intervals) overlapping with the coordinate range covered by each leaf. The generated index for each data collection allows for efficient access to the genomic records overlapping a query interval across all the tracks stored in the data collection.

Using these data indexes, the data collection tracks can then be efficiently accessed using the FILER Giggle-based query engine (see **Supplementary Methods**).

#### 2.2.5. Data access and availability

Data is added to the FILER periodically, with public versioned releases (data freezes) every six months. All data is physically stored in Amazon AWS (Amazon Web Services (AWS) - https://aws.amazon.com/) cloud. Access to the data is provided using FILER web server (https://lisanwanglab.org/FILER). Importantly, a local copy of FILER can be deployed on user-own servers or cloud computing instances (https://bitbucket.org/wanglab-upenn/FILER) to enable integration with existing or new analysis pipelines using the provided installation and data scripts.

## 3. Results

### 3.1. FILER contents

FILER contains functional genomics and annotation data characterizing various biological features of the human genome. **Figure 3** shows the distribution of functional genomic datasets integrated into FILER by data source, experimental data types, tissue/cell type categories, and biological sample types. FILER integrates functional genomic and annotation data across many primary data sources (**Figure 3a**) with the genomic/annotation records spanning a wide range of experimental assays/data types (**Figure 3b**). The data tracks in FILER are generated from a variety of biological sources (**Figure 3c**) and span a variety of tissue/cell type categories (**Figure 3d**) corresponding to >1,000 tissues/cell types. **Supplementary Figure S2** shows the distribution FILER genomic/annotation records across tissue/cell types categories, individual types of cells/tissues and data sources. Currently, FILER includes datasets across 14 human tissue and organ systems including cardiovascular (19.5%), nervous (15.9%), reproductive (8.9%), digestive (13.1%), stem cell (9.1%) and other categories.

**Figure 3.**
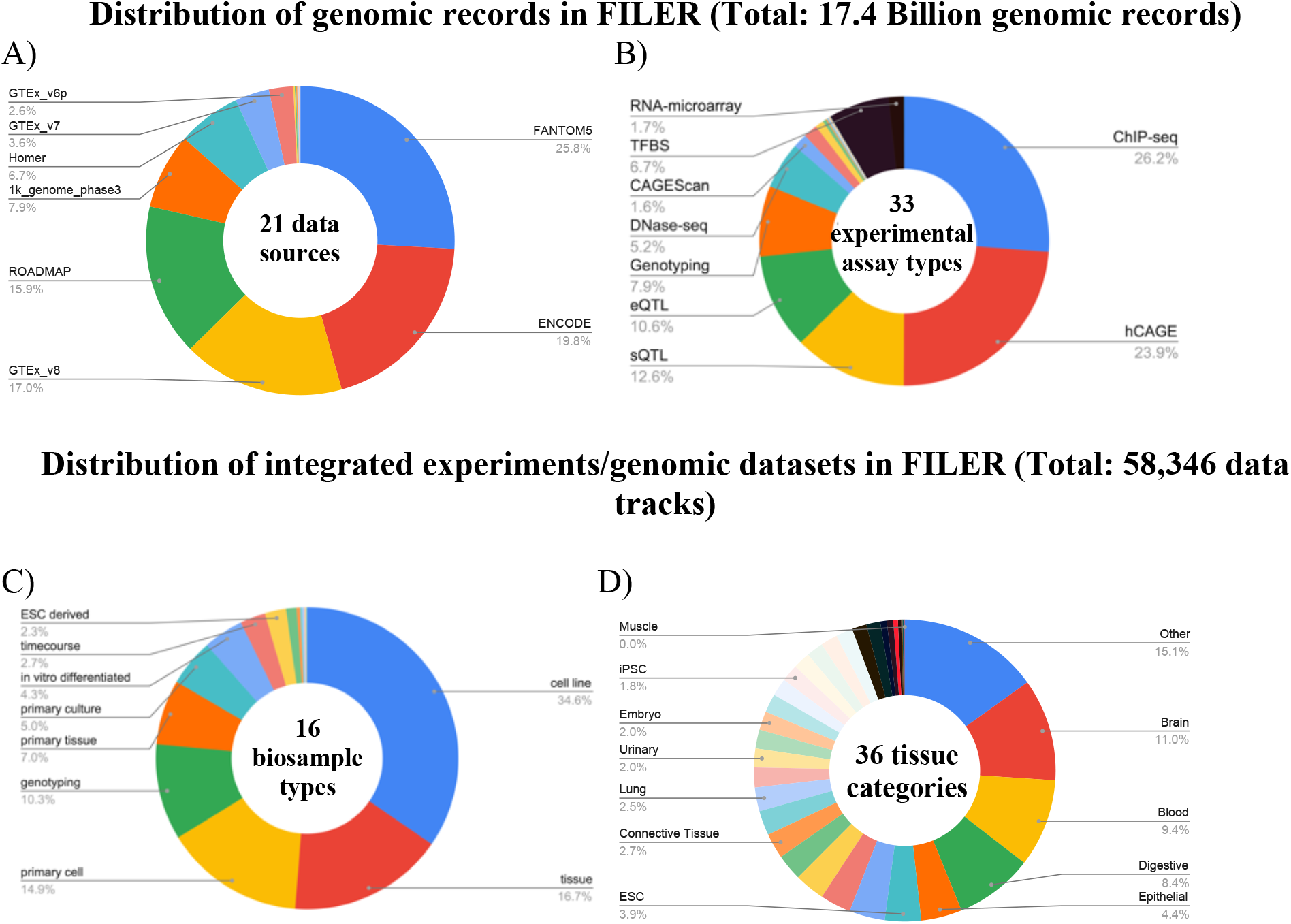
FILER data composition. A) Composition of functional genomic records by data source/consortium. B) Distribution of genomic records across experimental assays and data types. C) Distribution of datasets by biological source. D) Distribution of datasets across FILER tissue/cell type categories.

**Supplementary Table S7** provides a summary of the FILER datasets/records by the type of biological annotations and genomic features. FILER rich genomic and annotation data collection includes functional data capturing regulatory elements such as enhancers, transcription factor activity (transcription factor binding sites (TFBS) profiled by ChIP-seq (Park, 2009; Davis *et al.*, 2018; Dunham *et al.*, 2012) and computational analyses (Wang *et al.*, 2013; Heinz *et al.*, 2010)), quantitative trait loci (QTL) including expression QTL (eQTL) and splicing QTL (sQTL), chromatin state (ChIP-seq (Park, 2009; Davis *et al.*, 2018; Dunham *et al.*, 2012), DNase-seq (Song and Crawford, 2010), ATAC-seq (Buenrostro *et al.*, 2015)), gene transcription activity (transcription start sites (TSS) profiled by CAGE (Andersson *et al.*, 2014)), long-noncoding RNAs, small RNA loci (Kuksa *et al.*, 2019; Leung *et al.*, 2016), miRNA-mRNA interactions (Agarwal *et al.*, 2015), histone marks (ChIP-seq) (Park, 2009; Davis *et al.*, 2018; Dunham *et al.*, 2012; Kundaje *et al.*, 2015) and others (**Figure 3)**. In the case of replicated experiments, e.g., for histone and transcription factor ChIP-seq ENCODE experiments, in addition to peaks obtained for each replicate, more stringent/reproducible peak sets may be available including sets of replicated peaks, optimal or conservative IDR (irreproducible discovery rate) peaks sets (Landt *et al.*, 2012).

### 3.2. FILER data aggregation and integration

FILER data are derived by harmonizing and integrating functional genomics and annotation datasets (**Methods; Sections 2.1, 2.2**) across various primary data sources/consortiums (21 in FILER v1.0). **Supplementary Table S2** provides details of the primary data sources and experimental data types integrated into FILER.

FILER performs data integration at three levels: 1) individual dataset description (consistent meta information describing specifics of each of individual datasets); 2) grouping of datasets into broader categories across data sources (see **Methods**); and 3) data file formats (consistent, uniform file formats) (see **Methods**).

In particular, each individual FILER dataset is described by a consistent, standardized set of information attributes (metadata, >30 features; **Supplementary Table S4**), including biological sample, tissue/cell type, experimental protocol, library characteristics, along with file meta-information. This standard set of FILER attributes allows for consistent organization, presentation and retrieval of various datasets across data sources, biological conditions, and experimental data types (see **Section 3.3**; **Supplementary Figure S5**).

Further integration across data sources included in FILER is achieved by categorization of individual datasets into broader tissue/cell type, systems, and biological data categories (**Methods; Supplementary Methods**). Currently, FILER tracks span 36 tissue/cell type categories (e.g., see **Supplementary Figure S2** for a summary of data tracks by tissue/cell type category) and include data across 162 fine-level biological data categories (see **Supplementary Table S7** for a summary of biological data classification in FILER) and 14 system-level categories. This integrative categorization allows FILER to be used for viewing, filtering and aggregating data and query results across various tissue/cell types contexts and data sources (see, e.g., **Supplementary Figure S5**). Furthermore, all datasets integrated into FILER use consistent, uniform data file formats (BED-based) (**Methods;** see **Supplementary Table S6** for details of BED formats available in FILER) allowing for seamless integration, retrieval, and comparison of datasets across data sources, experimental data types (see **Sections 3.3** ‘FILER features’ and **3.4** ‘FILER example use cases’ for examples).

The current version of FILER includes 152 indexed data collections (**Methods**). **Supplementary Table S3** provides details on the data collections available in FILER.

Indexing of the individual data collections allows FILER to provide a scalable interface for querying (**Methods; Supplementary Methods**) genomic datasets by genomic coordinates or analysis and annotation of users’ own experimental data (**Section 3.3**).

FILER provides query/search functions (**Section 3.3**), allowing to easily integrate and query datasets using custom genetic and genomic analysis workflows such as INFERNO (Amlie-Wolf *et al.*, 2018; Kuksa *et al.*, 2020) (see **Section 3.4** for an example). The scalable interface of FILER allows analysis and annotation of user-generated experimental data (e.g., for a particular biological condition) with reference FILER datasets across various tissues/cell types and data sources (**Section 3.3**), facilitating research studies across different human diseases.

### 3.3. FILER web-portal features

FILER aims to provide a unified functional genomics and annotation resource to the scientific community. Currently, FILER allows the users to

i. quickly stage (identify and retrieve) relevant experimental datasets in specific tissue contexts for downstream analyses (**Supplementary Figure S5**);
ii. efficiently search and retrieve all genomic records (annotations and experimental features) across various data sources within a genomic region of interest or a set of genomic regions (**Supplementary Figure S3**);
iii. analyze and annotate user-provided experimental data using the reference datasets in FILER (**Figure 4**);
iv. deploy locally or on the cloud the entire FILER or a custom/selected subset of the FILER data.

**Figure 4.**
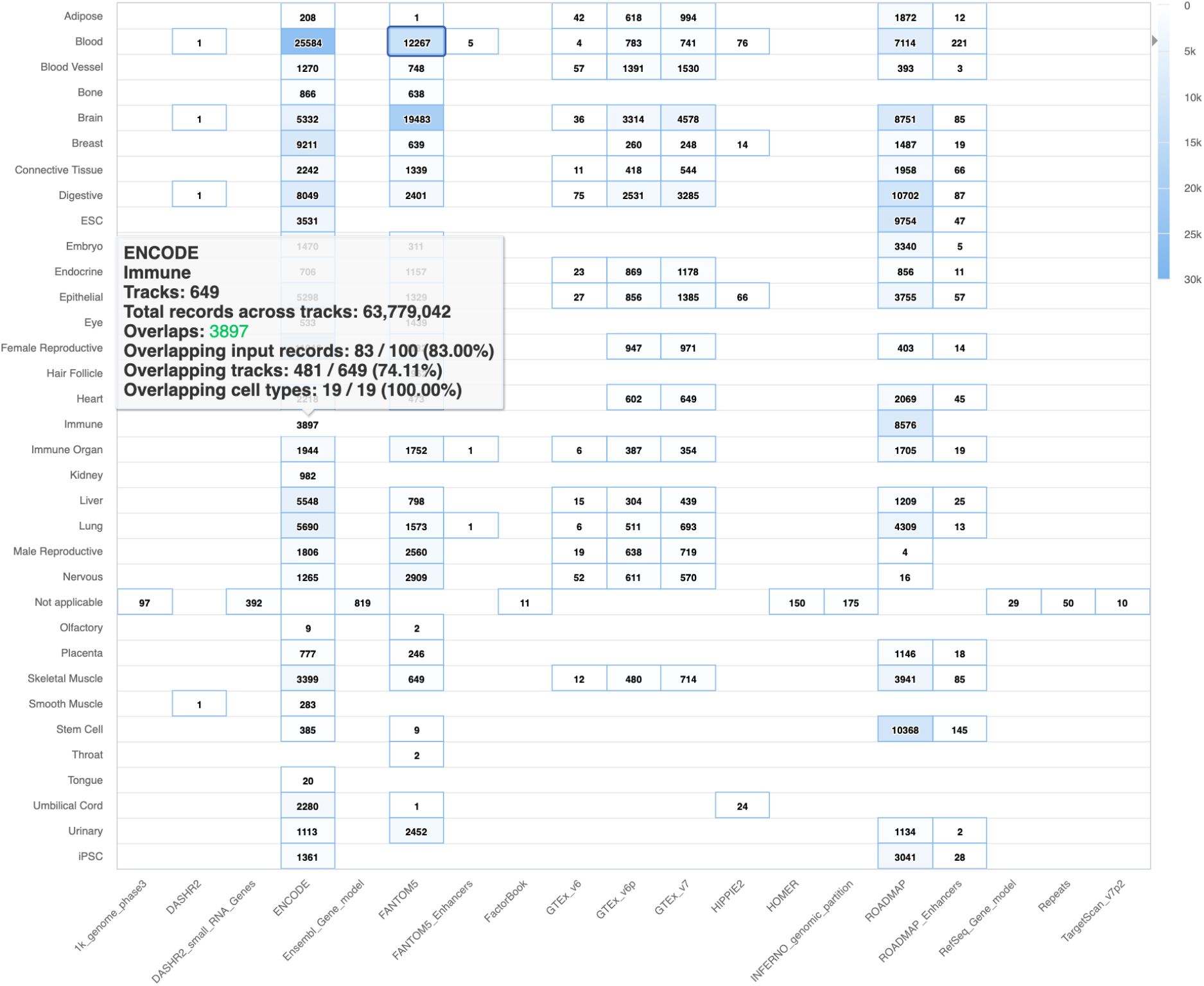
Example FILER output. Shown is an example of an interactive heatmap summarizing the distribution of overlaps for the ChIP-seq ENCODE input (ENCFF000AIA) across data sources (columns) and FILER tissue categories (rows). The interactive heatmap of the overlap results provided by FILER allows users to download results for any particular tissue category and data source. For each tissue category and data source, FILER provides a brief summary of the overlaps indicated the number of overlaps, the number of overlapping tracks, and the number of individual cell/tissue types. Similarly, FILER also provides an interactive heatmap (not shown in the figure) for experimental assays (columns) and tissue categories (rows).

**Supplementary Table S1** shows a comparison of the FILER features with currently existing tools and web servers.

#### 3.3.1. Search

FILER allows users to search for relevant genomic datasets based on their properties (meta-information). The genomic records (genomic features and annotations) across datasets can also be searched using FILER based on a) genomic coordinates of the region of interest or b) a set of coordinates for the genomic loci, e.g., experimentally-derived regions. Next, we describe the search functionality provided by FILER in more detail.

**Search by genomic interval:** Search by genomic interval (**Supplementary Figure 3a**) in FILER allows users to locate genomics data tracks and features overlapping with the given genomic coordinates. All the overlapping genomic features, data tracks, and their associated meta-data will be returned (see **Supplementary Table S9** for an example of annotated overlap BED file; descriptions of the file columns are provided in **Supplementary Table S4**).

**Analyze and annotate user’s experimental data:** As shown in **Supplementary Figure S3**, the experimental data is provided in BED format. The user BED file can either be uploaded to the FILER web server (‘Upload your data’ tab), or provided through a web-accessible URL (‘Analyze your own data’ tab).

**Browse and search genomic datasets by data source, tissue/cell type, assay, data type and other properties:** FILER allows users to locate relevant data tracks by a given metadata attribute or a combination of attributes (see **Supplementary Figure S5; Section 3.4, Use Case 1** for an example). The selected data tracks matching the search parameters can be then downloaded including the processed, as well as the original (raw) datasets along with their associated meta-data.

#### 3.3.2. Analysis and annotation of user-provided experimental data

FILER allows users to locate all data tracks and genomic features overlapping user-provided experimental intervals. The experimental intervals in the user-provided BED file are compared with all FILER datasets and all overlapping genomic features, data tracks and meta-data are reported (**Supplementary Figure S4**). The full set of results are available for download in the compressed (zip) format. The main annotated BED file containing all overlaps and FILER metadata is also available for download. FILER outputs a set of results tables and figures generated by matching the set of genomic loci in the user/input experiment against sets of genomic loci in each tissue and cell type in the reference FILER datasets (**Supplementary Figure S4**). As FILER performs this head-to-head comparison of the genomic loci in the input BED file with all functional genomics datasets in FILER, this functionality allows users to annotate and compare easily any experimental data of their own interest against all the FILER reference datasets.

### 3.4. FILER example use cases

Broad tissue/cell type coverage and a wide range of experimental data types available in FILER enable systematic analyses of genome-wide studies such as experimentally-derived ChIP-seq or RNA-seq genomic intervals or association signals observed in GWASs.

We next describe example use cases for FILER, including 1) using a FILER web server for identification and retrieval of relevant experimental datasets (**Use case 1**), 2) using custom/user experimental data with FILER webserver (**Use case 2**), and 3) integration into high-throughput analysis workflows (**Use case 3**) using FILER installation and data command-line scripts.

1. **Use case 1**: relevant experimental data retrieval/data staging. FILER can be used to identify and download relevant datasets using various criteria, including a particular human tissue context (tissue category), a particular experimental data type (assay), or a specific data source. As shown in **Supplementary Figure S5**, via the web-interface, data selectors allow users to choose specific data of interest. All the data tracks matching the selection criteria can be download in bulk or individually, including the associated track meta-information. Alternatively, the command line interface provided by FILER (see **Supplementary Methods**, ‘Deploying FILER data’ section) allows users to deploy the relevant data subset of interest locally or on a cloud using the FILER meta-information table. The metadata will serve as a guide to download individual data tracks, re-create FILER data structure, and generate data indexes (see also **Use case 3** for an example of integrating FILER data with a high-throughput workflow; **Supplementary Methods**, ‘Deploying FILER data’).
2. **Use case 2**: analyzing custom experimental data. FILER can be used for analysis of users’ own experimental data (e.g., experimentally-derived genomic intervals). In particular, FILER can be used to annotate custom genomic data from the user experiment, such as ChIP-seq peaks, RNA-seq small RNA peaks, or ATAC-seq open chromatin regions. Using the provided BED file with experimental data as input, FILER will generate an annotated BED file containing all genomic records overlapping with the user-provided experimental intervals (available for download under ‘Annotated overlaps’ subsection). The annotated overlap file will contain overlap records consisting of the three sections: 1) all the data fields in the user input, 2) overlapping FILER data track information, and 3) the associated FILER track meta-information. Along with the main annotated overlap file in BED format, FILER will generate a report detailing the distribution of overlapping genomic records across tissue categories, data sources, and experimental data types. Additionally, overlapping genomic records corresponding to a particular tissue category/data source, or tissue category/experimental assay are available for individual download and further analysis. In **Figure 4**, we showed the FILER results via the web-portal when analyzed using a sample of ChIP-seq ENCODE data (ENCODE accession ENCFF000AIA; 92,008 ChIP-seq peaks) as an example. This interactive heatmap allows users to download results in BED format for any specific tissue category and data source.
3. **Use case 3**: integration with a genetic and genomic analyses workflow. **Figure 5** shows an example of integrating FILER with a custom aggregation and analysis pipeline. In this example, FILER data is deployed on Amazon AWS cloud (see **Supplementary Methods**, ‘Deploying FILER data’ for details) and the SparkINFERNO (Kuksa *et al.*, 2020) pipeline for non-coding variant analysis is using FILER datasets and FILER genomic query engine to characterize the input GWAS genetic variants. The annotated overlap data output spans across several biological data types, including enhancer, transcription factor (TF) binding, and open chromatin genomic features.

**Figure 5.**
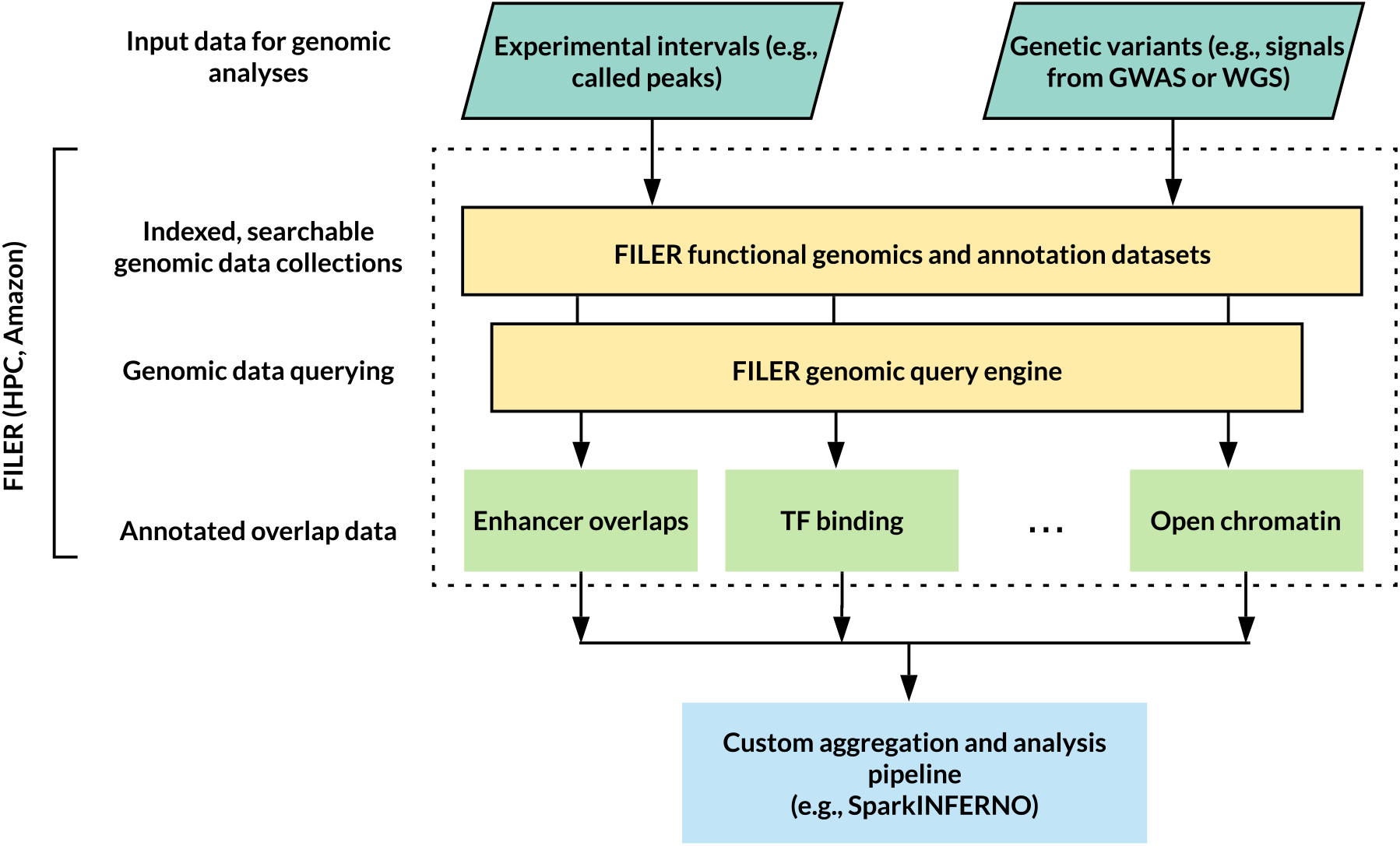
Example of integrating FILER (middle part of the figure) with a custom aggregation and analysis workflow (bottom of the figure), SparkINFERNO (Kuksa *et al.*, 2020) for high-throughput non-coding variant analysis (inputs at the top).

To demonstrate the utility of FILER in analyzing and characterizing genomic regions or features of interest, we used FILER together with SparkINFERNO pipeline (Kuksa *et al.*, 2020) for functional characterization of non-coding variants in IBD GWAS (Liu *et al.*, 2015). As part of the SparkINFERNO pipeline, all genome-wide significant GWAS variants (p-value<5 × 10^-8^) were used to obtain a set of candidate, potentially causal IBD variants by linkage disequilibrium (LD)-based pruning and expansion of the genome-wide significant GWAS SNPs. All the resulting 16,694 candidate variants were overlapped against GRCh37/hg19 FILER tracks (22,123 tracks, 7.1 Billion genomic records). In total, over 40 million overlaps with FILER genomic records were found. The overlap of these variants across functional genomic categories in different tissue types is shown in **Supplementary Figure S6**. As can be seen from the figure, overlap of IBD variants with the FILER data highlighted tissues that are likely important for IBD disease etiology, including blood, immune and digestive categories (Amlie-Wolf *et al.*, 2018; Liu *et al.*, 2015).

### 3.5. FILER deployment

FILER is available in two different ways:

1. FILER data is accessible through the web server (https://lisanwanglab.org/FILER). It allows users to search and retrieve relevant data, retrieve genomic features and annotation for genomic regions of interest, and analyze and annotate user’s experimental data with the reference FILER datasets.
2. FILER can be deployed locally on a server or cluster (https://bitbucket.org/wanglab-upenn/FILER). This not only allows users to access harmonized functional genomics and annotation data stored in FILER, but also integrate FILER into custom analysis pipelines. The scalable, genomic indexing-based interface allows to efficiently access FILER large-scale functional genomics data collection and use it in custom analyses.

FILER code repository (https://bitbucket.org/wanglab-upenn/FILER) provides installation and data preprocessing scripts to download, prepare metadata, and index FILER data.

**Supplementary Methods**(‘Deploying FILER data’) provides details on how to install and use FILER locally.

### 3.6. FILER data access and querying API

All FILER data and metadata can be accessed programmatically using command-line interface. Command-line scripts for accessing/querying FILER data are available in the FILER code repository (https://bitbucket.org/wanglab-upenn/FILER). Individual track data and track metadata can be accessed, e.g., using the provided `get_region_data.sh` and `get_metadata.sh` scripts. Tracks in FILER can also be queried by a genomic interval of interest (e.g., `get_overlapping_tracks_by_coord.sh` script). For details on usage and example commands please refer to the README and help for individual scripts.

## 4. Discussion

Analyses of the results from GWASs and biological experiments require using external functional data as evidence for further interpretation, characterization and discovery. However, there is currently no single resource that provides unified access to a harmonized collection of such functional genomics and annotation data. This complicates the search and use of relevant functional genomic data, as well as the comparison and aggregation of these heterogeneous datasets for the analyses. More importantly, without a unified, harmonized data resource it is difficult to use these valuable resources in various genomic and genetic pipelines. We envision that the large-scale, harmonized FILER genomic and annotation data collection will facilitate the downstream genetic and genomic analyses, including but not limited to studying GWAS signals, via system biology, causal gene and other analyses. This allows researchers to focus on the creative analysis tasks, genomic analyses and discoveries rather than data collection and cleaning.

FILER uniquely provides an integrated and extensible repository of harmonized functional genomics and annotation data allowing for efficient and seamless retrieval, analysis and comparison across data sources, biological conditions, tissues/cell types, and experimental data types. Using provided deployment and data interface, FILER allows integration with the existing or new analytical workflows. In addition to the web-based access, FILER can be installed on a local server, high-performance computing (HPC) cluster or cloud computing instances (see **Section 3.5; Supplementary Methods**).

Broad tissue/cell and experimental data type coverage in FILER enables systematic analyses of genome-wide experiments such as ChIP-seq or ATAC-seq, or genome-wide analyses of association signals observed in GWASs. Moreover, the modular, data collection-based FILER data architecture allows additional analysis-specific/user data or new data sources to be easily added without affecting other datasets/data collections.

Current limitations of the FILER web server include the usage of a single multi-core server with local data storage. Implementation based on Apache Spark (Zaharia *et al.*, 2016) or similar distributed computing framework and distributed data storage would allow further scalability in terms of both the data size and the search and processing speed.

Future developments for FILER will include 1) broadening of tissue/cell type coverage, 2) adding and expanding experimental data types including chromatin interaction/3D genome organization data, gene and RNA expression, and single-cell experiments, 3) adding disease-specific datasets, 4) variant-level annotations, 5) visualization, interactive display and exploration of functional genomic and annotation datasets.

## Supporting information

Supplementary Table S2. Summary of integrated data sources

Supplementary Table S3. FILER data collections

Supplementary Table S4. FILER metadata schema description

Supplementary Table S5. Schema matching for integrated data sources

Supplementary Table S6. Data file formats/schemas for data tracks in the FILER database

Supplementary Table S7. Summary of FILER data across biological annotation/data categories

Supplementary Table S8. FILER metadata table

Supplementary Table S9. Example of FILER annotated overlaps

Supplementary Note S1. FILER data collection and preparation

## 5 Data availability

All harmonized genomic datasets and the corresponding annotation meta-data are freely available from the FILER website https://lisanwanglab.org/FILER. An entire FILER database or a selected data subset can be deployed on local server, cloud or high-performance computing (HPC) clusters using installation and distribution scripts provided at https://bitbucket.org/wanglab-upenn/FILER.

## 6. Acknowledgment

We thank Mitchell Tang for setting up, collecting and organizing the initial collection of genomic and annotation tracks. We also thank the members of Wang Lab for their comments and insightful discussions.

## Supplementary Figures and Tables

**Supplementary Figure S1**. FILER highly efficient genomic search and retrieval

FILER_supplementary.pdf

**Supplementary Figure S2**. Distribution of genomic/annotation records across FILER tissue/cell categories

FILER_supplementary.pdf

**Supplementary Figure S3**. FILER search interface

FILER_supplementary.pdf

**Supplementary Figure S4**. Analysis and annotation of the user-provided experimental data using FILER

FILER_supplementary.pdf

**Supplementary Figure S5**. Retrieving genomic and annotation datasets of interest using FILER

FILER_supplementary.pdf

**Supplementary Figure S6**. Characterization of top genome-wide significant IBD variants.

FILER_supplementary.pdf

**Supplementary Table S1.** Summary of features for FILER and other similar tools

FILER_supplementary.pdf

**Supplementary Table S2.** Summary of integrated data sources

FILER_supplementary_Table_S2_data_source_summary_with_ref.xlsx

**Supplementary Table S3.** FILER data collections

FILER_supplementary_Table_S3_data_collections_summary.xlsx

**Supplementary Table S4.** FILER metadata schema description

FILER_supplementary_Table_S4_metadata_description.xlsx

**Supplementary Table S5.** Schema matching for integrated data sources

FILER_supplementary_Table_S5_schema_matching.xlsx

**Supplementary Table S6.** Data file formats/schemas for data tracks in the FILER database

FILER_supplementary_Table_S6_file_formats.xlsx

**Supplementary Table S7.** Summary of FILER data across biological annotation/data categories

FILER_supplementary_Table_S7_biological_data_category_summary.xlsx

**Supplementary Table S8.** FILER metadata table

FILER_supplementary_Table_S8_metadata.zip

**Supplementary Table S9**. Example of FILER annotated overlaps

FILER_supplementary_Table_S9_annotated_overlaps_example.xlsx

**Supplementary Note S1.** FILER data collection and preparation

FILER_supplementary_Note_S1_data_collection_and_preparation.docx

