## Supplementary information for "FILER: large-scale, harmonized FunctIonaL gEnomics Repository"

Mailing address: Richards Building, D101, 3700 Hamilton Walk, Philadelphia, PA 19104

Yuk Yee Leung, PhD

Mailing address: Richards Building, D102, 3700 Hamilton Walk, Philadelphia, PA 19104

### Supplementary information

### 1. Supplementary Material and methods

#### 1.1. Data collection and curation

FILER integrates data from multiple genomic databases including ENCODE (Davis *et al.*, 2018; Dunham *et al.*, 2012), FANTOM5 (Andersson *et al.*, 2014), ROADMAP Epigenomics (Roadmap Epigenomics Consortium *et al.*, 2015), GTEx (Aguet *et al.*, 2020, 2017), DASHR (Kuksa *et al.*, 2019; Leung *et al.*, 2016) (see **Supplementary Table S2** for details of data sources available in FILER). ENCODE data in FILER includes datasets from 19 experimental assays including ChIP-seq (Park, 2009; Davis *et al.*, 2018; Dunham *et al.*, 2012), DNase-seq (Song and Crawford, 2010), ATAC-seq (Buenrostro *et al.*, 2015). FILER includes the genomic data based on GRCh37/hg19 and GRCh38/hg38 human reference genomes. Genomic datasets that were only available for GRCh37/hg19 were lifted-over (Kuhn *et al.*, 2013) to the GRCh38/hg38 build and were also included in FILER. Genotype-Tissue Expression (GTEx) data includes both expression quantitative trait loci (eQTL) and splicing quantitative trait loci (sQTL) data (GTEx v6p, GTEx v7 and GTEx v8 data are available in FILER). Release 9 of the ROADMAP epigenomic data for 183 biological samples is included in FILER. Users can also access ROADMAP ChromHMM enhancers (Ernst and Kellis, 2017, 2012; Roadmap Epigenomics Consortium *et al.*, 2015). FILER also includes the expressed enhancer tracks in addition to the CAGE data from FANTOM5 (Andersson *et al.*, 2014). All of the FANTOM5 data are lifted over to GRCh38/hg38 genome build in FILER. Transcription factor binding and motif data from Factorbook (Wang *et al.*, 2013) and Homer (Heinz *et al.*, 2010) are available in both GRCh37/hg19 and GRCh38/hg38 genome builds. Annotations for microRNA targets from TargetScan (Agarwal *et al.*, 2015) and small human noncoding RNAs annotations from DASHR (Kuksa *et al.*, 2019; Leung *et al.*, 2016) are included. Ensembl (Yates *et al.*, 2020) and RefSeq (O'Leary *et al.*, 2016) gene models, repeat region annotations (Tempel, 2012) and reference genome are available in both GRCh37/hg19 and GRCh38/hg38 genome builds.

All datasets downloaded from the multiple sources are pre-processed and converted into the standard BED-based (Kent *et al.*, 2002) format for consistency (see **Supplementary Note S1** 'FILER documentation' for details).

All tracks included in FILER were annotated with an extensive meta-information including original track information, data source, experimental assay, biological source, file meta-data (see **Supplementary Table S4** for details on the meta-information provided). All tracks in FILER were categorized by biological feature and data type based on a combination of attributes including the experimental assay, experimental target (e.g., antibody used; if available) and genomic record and feature type (e.g., peaks, eQTL, transcription start sites, and others).

To provide efficient access to all tracks, FILER stores all tracks as individually-indexed data collections (see **Supplementary Table S3** for a list of data collections available in FILER). To provide a modular and extensible data storage architecture, tracks are stored in a separate directory for each data source. Each data source is then further organized by experimental assay, data type, and genomic build into sub-directories. The last level of the directories corresponds to individual data collections. Each data collection thus holds tracks of the same type and data format (see **Supplementary Table S3** for the details of data organization and FILER data collections). Each data collection is then indexed based on Giggle genomic indexing (Layer *et al.*, 2018). The index provides efficient access/querying based on a single genomic region or a collection of genomic regions of interests (see **Supplementary Figure S1** for an example). FILER is based a customized version of Giggle (available at https://github.com/pkuksa/FILER_giggle) providing indexing, querying/genomic search and output of the results in the BED-based format.

FILER is continuously updated, with data freezes every 6 months. New release information will be updated at https://lisanwanglab.org/FILER.

**Supplementary Note S1** ('FILER data collection and preparation') includes more detailed information on data collection for each of the data sources, meta-information extraction/generation, and raw track data pre-processing.

Each data-source specific metadata and schema were matched across data sources (**Supplementary Table S5** provides the details of the schema matching) to generate consistent meta-data descriptions for each of the FILER tracks.

#### 1.2. Processing and quality control

Each individual dataset was converted from its original format (e.g., GFF (GFF2 - http://gmod.org/wiki/GFF2), bigBed (Kent *et al.*, 2010) into the standard BED-based format to enable indexing and efficient querying. Genomic coordinates were converted when needed to the BED-based 0-based format. Chromosome names were standardized to use the consistent chr[0-9XYM] notation. Lifted data tracks included the lifted genomic coordinates (GRCh38/hg38) as the first columns followed by the all of the original fields in the GRCh37/hg19 data file. All BED files were compressed using `bgzip` (Li, 2011) to reduce storage footprint and enable both Giggle-based indexing of set of tracks and `tabix` (Li, 2011) indexing of individual tracks.

#### 1.3. Tissue categorization

As FILER integrates data across several data sources and consortiums, the different tissue/cell type categorizations from the integrated data sources had to be combined into a single, consistent reference for tissue/cell type categorization to allow for integration of datasets across data sources. The steps described below outline the tissue/cell type categorization process for NIH Roadmap Epigenomics, FANTOM 5, ENCODE, DASHR and all other data sources (Ernst and Kellis, 2012; Roadmap Epigenomics Consortium *et al.*, 2015; Ernst and Kellis, 2017; Heinz *et al.*, 2010; Andersson *et al.*, 2014).

First, the set of standard tissue/cell type categories used for integration included 36 tissues: Adipose, Blood, Blood Vessel, Bone, Brain, Breast, Circulating Cell, Connective Tissue, Digestive, Embryo, Endocrine, Epithelial, ESC, Eye, Female Reproductive, Hair Follicle, Heart, Immune, Immune Organ, iPSC, Kidney, Liver, Lung, Male Reproductive, Muscle, Nervous, Olfactory, Placenta, Skeletal Muscle, Smooth Muscle, Stem Cell, Throat, Tongue, Tonsils, Umbilical Cord, Urinary.

All unique tissues/cell types in FILER were matched to one of the 36 categories. If the more broader tissue category was provided by the data source in the description of a dataset, this data-source tissue category was used for assigning the genomic dataset to the tissue/cell type category. When no information on the tissue context or cell type category was provided by the data source, the mapping of individual tissues/cell types to the tissue/cell type category was performed using BTO, CLO, CL and other ontologies (Gremse *et al.*, 2011; Schomburg *et al.*, 2013; Sarntivijai *et al.*, 2014; Diehl *et al.*, 2016; Malone *et al.*, 2010; Smith *et al.*, 2007). The tissue/cell type category was decided by tracing the specific cell type/cell line classifications provided in ontologies from the leaf nodes (specific cell type/cell line) to more general/more broad classifications represented by parent and other ancestor nodes. Extra care was taken to decide on cell types/cell lines/tissues that matched multiple tissue categories. For example, a category name such as “thalamus” would fall into both the nervous and brain categories in the mapping. The most specific (e.g., tissue of origin) category was chosen in most cases. In this example, “brain” was chosen as the final category.

An example of the cell type/tissue categorization process s shown below.

| Example of the tissue mapping process for Monocyte cell |
|---|
| 1. Adult stem cell -> hematopoietic stem cell -> myeloid progenitor cell -> marrow cell -> mononuclear phagocyte -> Monocyte |
| 2. Hematopoietic system -> **blood** -> blast cell -> hematopoietic **stem cell** -> hematopoietic progenitor cell -> myeloid progenitor cell -> marrow cell -> mononuclear phagocyte -> Monocyte |
| 3. Hematopoietic cell -> leukocyte -> Monocyte |
| 4. Hematopoietic cell -> marrow cell -> mononuclear phagocyte -> Monocyte |
| 5. Phagocyte -> mononuclear phagocyte -> Monocyte |

**The names that match one of the 36 tissues are highlighted

The tissue-base categorization has been applied to all the FILER data sources (Roadmap, Fantom 5, ENCODE, GTEx, DASHR and others; see **Supplementary Table S2** for the list of data sources integrated into FILER). The mapping for each unique cell type/tissue/cell line was summarized in a spreadsheet to avoid duplicates.

Overall, FILER contains comprehensive mapping of >1,100 tissues/cell types into 36 tissue categories (see FILER metadata in **Supplementary Table S8**). The tissue-based categorization allows users to integrate, retrieve and compare datasets from the various data sources integrated into FILER.

#### 1.4. Deploying FILER data

To create a local copy of the entire FILER (or a particular FILER data source/subset) for use with custom analysis pipelines, the FILER code repository (https://bitbucket.org/wanglab-upenn/FILER) provides an easy-to-use installation/deployment script (`install_annot.sh`). The installation script uses as input a metadata file template containing information on all of the data tracks to be deployed. The data installation is performed in two stages:
1) download FILER tracks referenced in the input metadata file and re-create FILER directory structure under the specified local/target directory, and
2) index individual data tracks using `tabix` (for all individual tracks found in the metadata file) and index all data collections using a customized version of `Giggle` (https://github.com/pkuksa/FILER_giggle).
The two steps are performed for all the data collections found in the input FILER metadata file.

For example, to install the latest GRCh38/hg38 FILER tracks into a local `FILER/` directory, the following command can be used:

```
bash install_annot.sh FILER
https://tf.lisanwanglab.org/GADB/metadata/filer.latest.hg38.template
```

Further examples of downloading a selected subset of FILER tracks (e.g., only ChIP-seq ENCODE, or tissue-specific enhancer tracks) are provided in the documentation for the FILER code repository (https://bitbucket.org/wanglab-upenn/FILER).

The FILER data can be stored in any local directory with the sufficient space on the target machine/server (e.g., /local/path/FILER), however since the absolute paths for data files are used to create the Giggle indexes, the local FILER data directory should not be moved to another place after indexing, otherwise re-indexing will be required.

Using the provided installation script with a custom metadata template allows to customize the set of FILER data tracks that will be downloaded, placed in the corresponding directories and indexed. A custom data template allows to create analysis-specific functional and annotation data collections. User data can be specified in the custom metadata template to further tailor the set of data sets to the data of interest for the analyses performed. Additional details and documentation on deploying and staging a custom subset of the FILER data is provided at the FILER code repository (https://bitbucket.org/wanglab-upenn/FILER).

### 2. Supplementary Figures and Tables

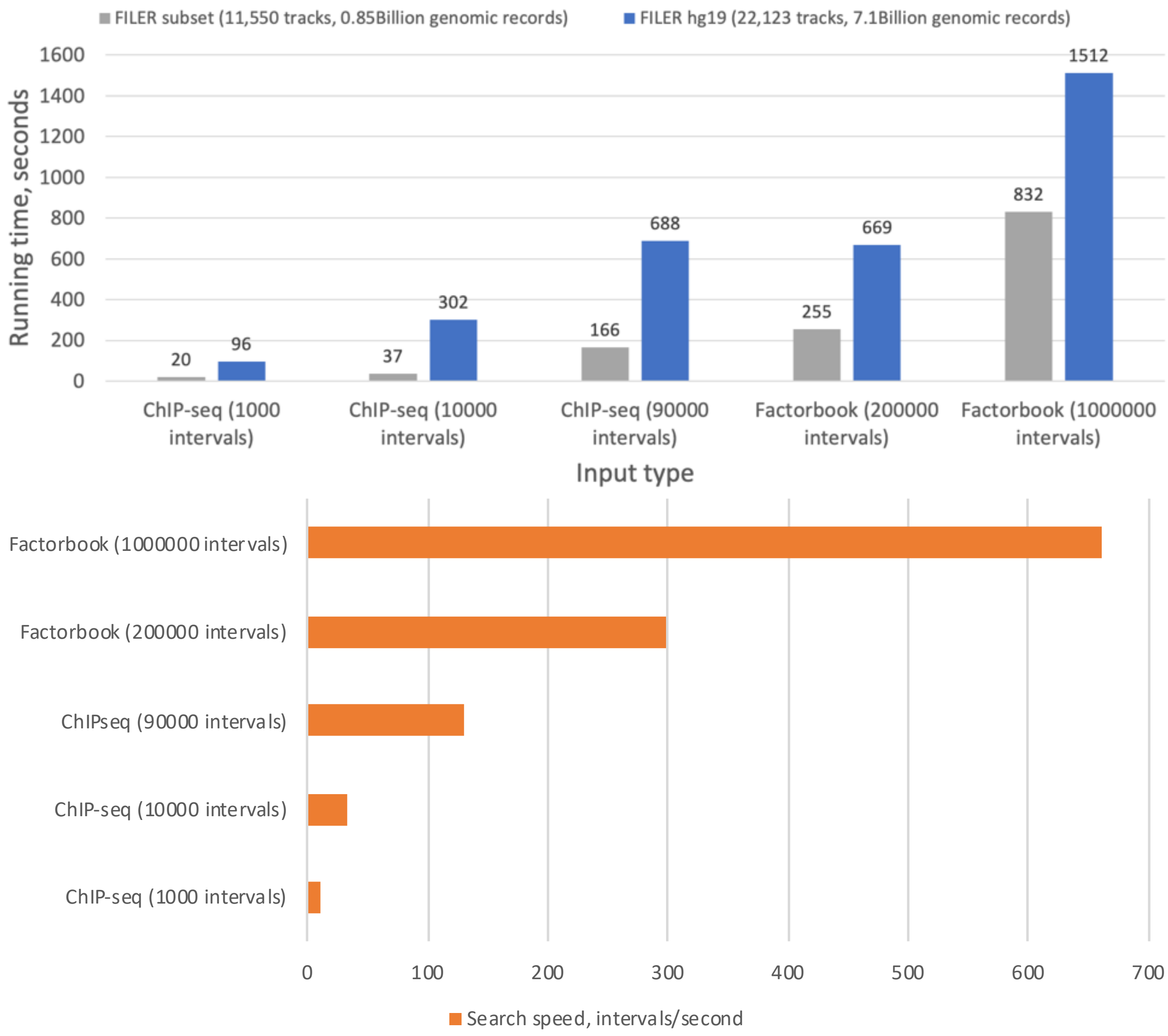


**Supplementary Figure S1**. FILER highly efficient genomic search and retrieval. **A)** Shown is the running time (in seconds) as a function of the input size and type (1,000 to 1,000,000 query intervals, ChIP-seq and TFBS data) and the amount of reference datasets (all hg19 FILER tracks with 7.1 Billion records and a ~10% subsample with 0.8 Billion records). Running time is measured on a AWS m5.4xlarge instance. **B)** Shown is the search speed (as the number of input intervals processed per second) using Factorbook transcription factor binding motif sites and ENCODE ChIP-seq peaks data against all hg19 FILER tracks.

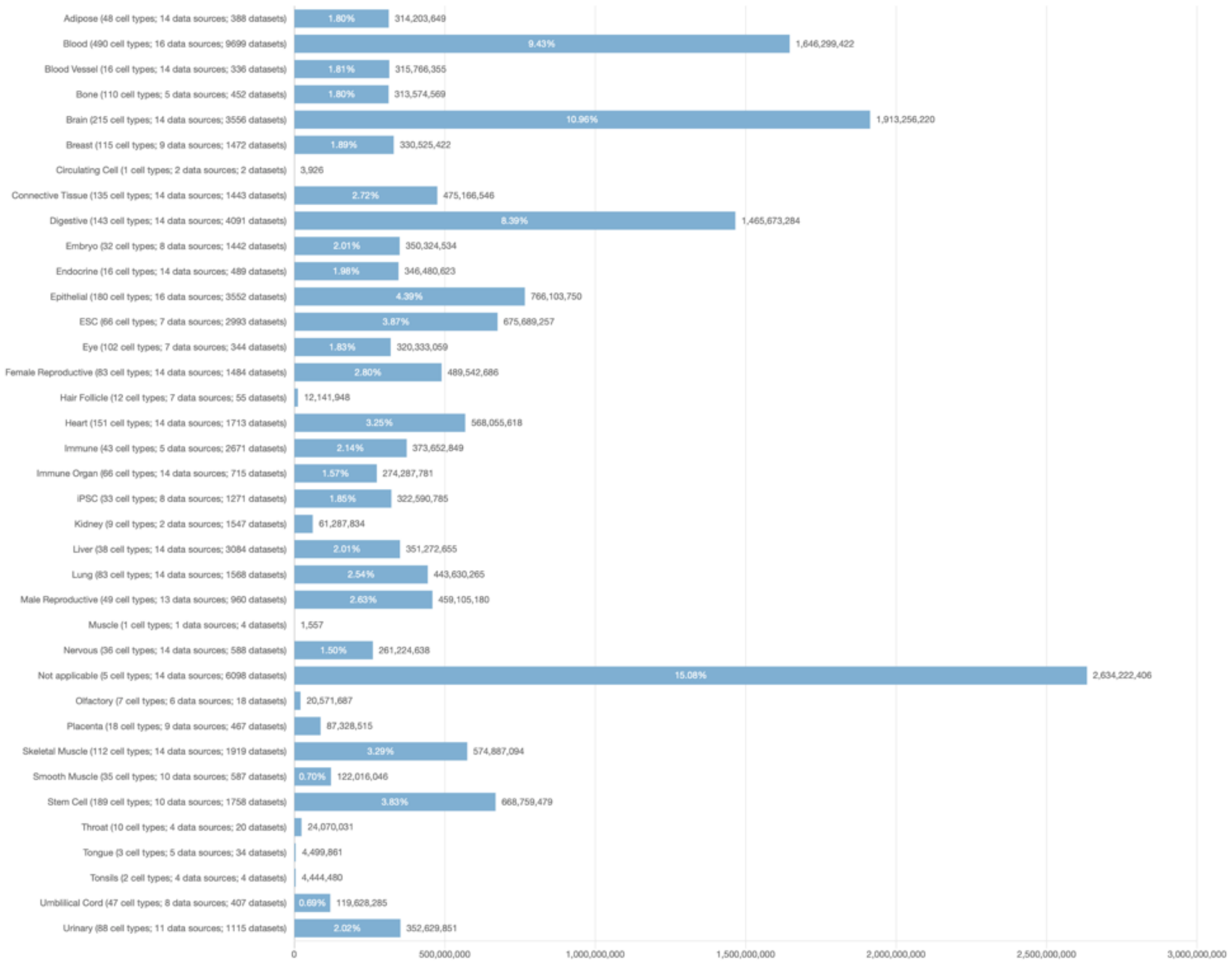


**Supplementary Figure S2**. Distribution of genomic/annotation records across FILER tissue/cell categories. We show for each tissue/cell type category, the number of records, the number of individual cell/tissue types corresponding to the tissue/cell type category, as well the number of contributing data sources and datasets.

**A)**

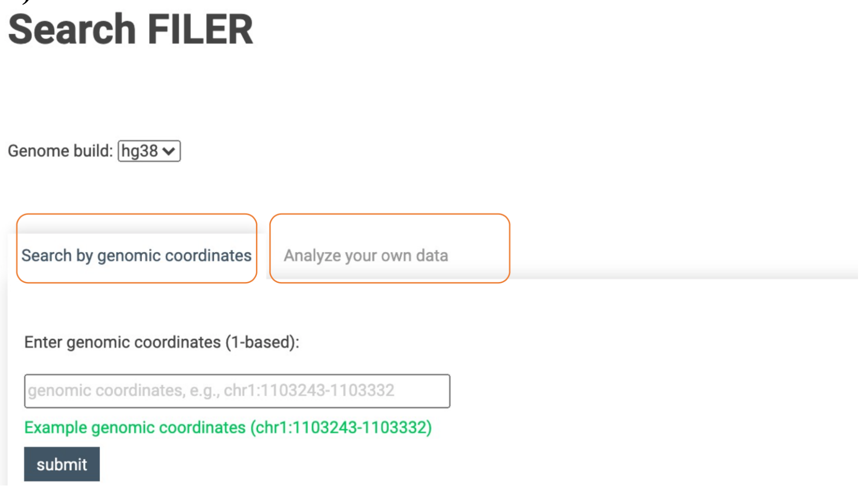


**B)**

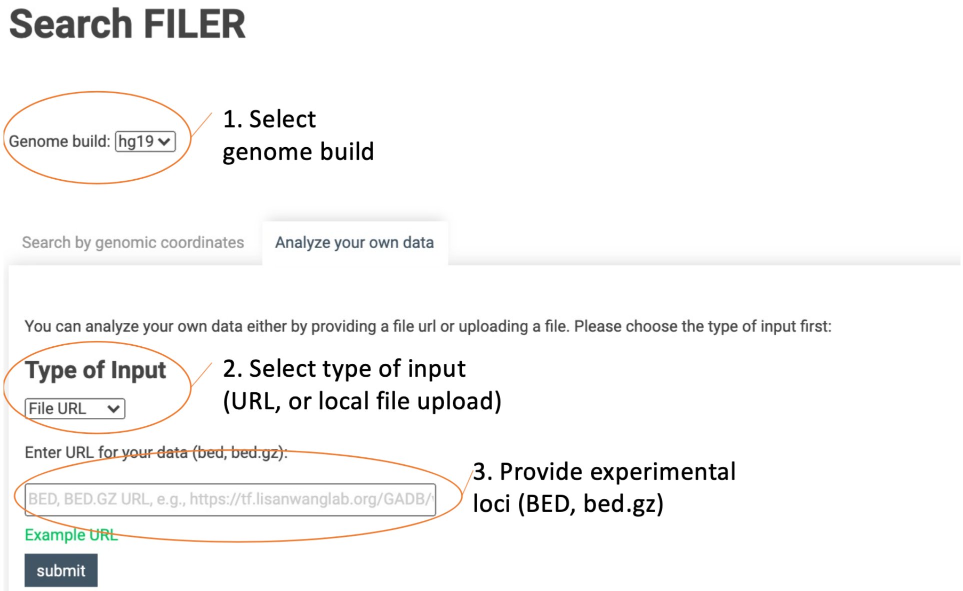


**Supplementary Figure S3**. FILER search interface. FILER provides an easy interface to analyze and annotate user data against reference FILER datasets/tissues without the need for installation. **A)** FILER data can be queried for any genomic region of interest ('Search by genomic coordinates') or using a set of experimental genomic loci (BED file) ('Analyze your own data'). Orange rectangles highlight the search/query modes available to users. **B)** Analysis of user's own data. Provided experimental loci (in plain BED or in the compressed bed.gz format) will be overlapped with FILER datasets. Results will be available for immediate download and will be kept for one week.

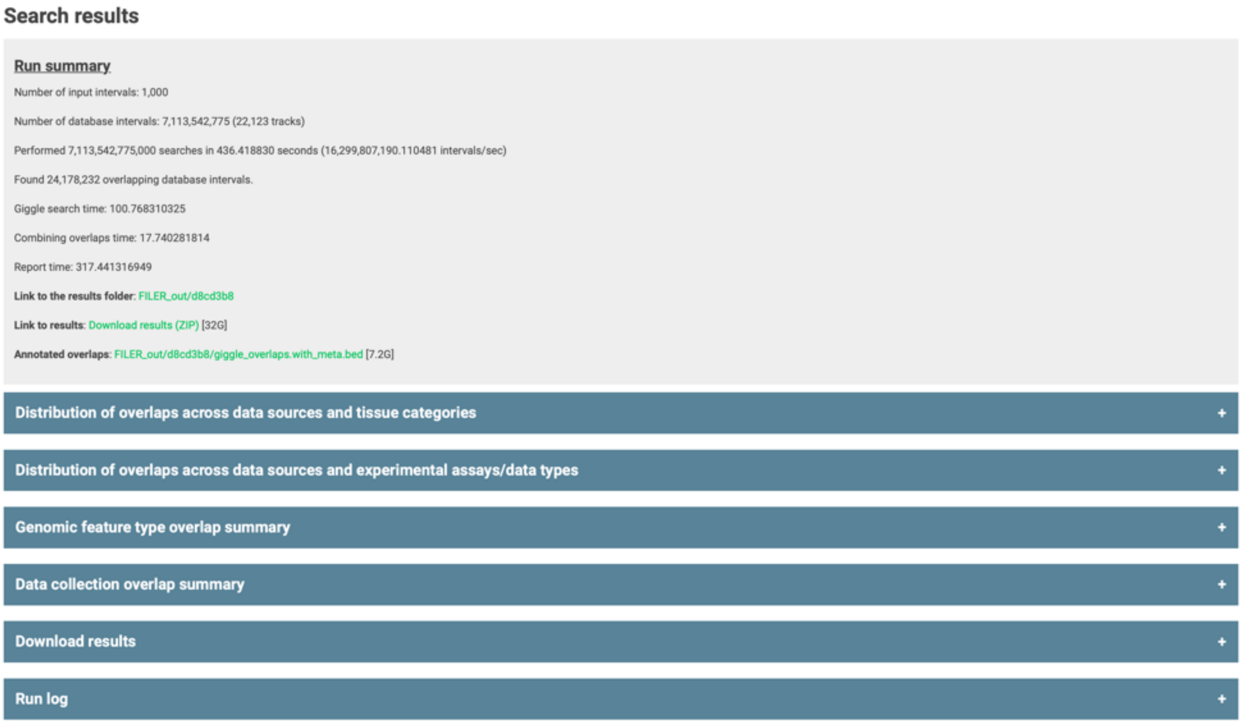


**Supplementary Figure S4**. Analysis and annotation of the user-provided experimental data using FILER. User-provided experimental loci (genomic intervals) (**Figure 4B**) are used to query FILER genomic and annotations tracks. FILER returns 1) a run summary containing search and overlap statistics and links to results, and 2) a set of overlap summaries across tissue categories, data sources, experimental assays, genomic features. Annotated overlap results include input data, overlap information and associated FILER metadata (annotations) for each of overlapping tracks. Annotated overlap results are available for download 1) as a single BED file containing all overlaps found in FILER (available under ‘Annotated overlaps’ subsection), 2) as separate BED files for each of tissue category/data source, tissue category/experimental assay pairs (available under ‘Distribution of overlaps’ subsections), and 3) for each of the genomic feature types (available under ‘Genomic feature type overlap summary’ subsection).

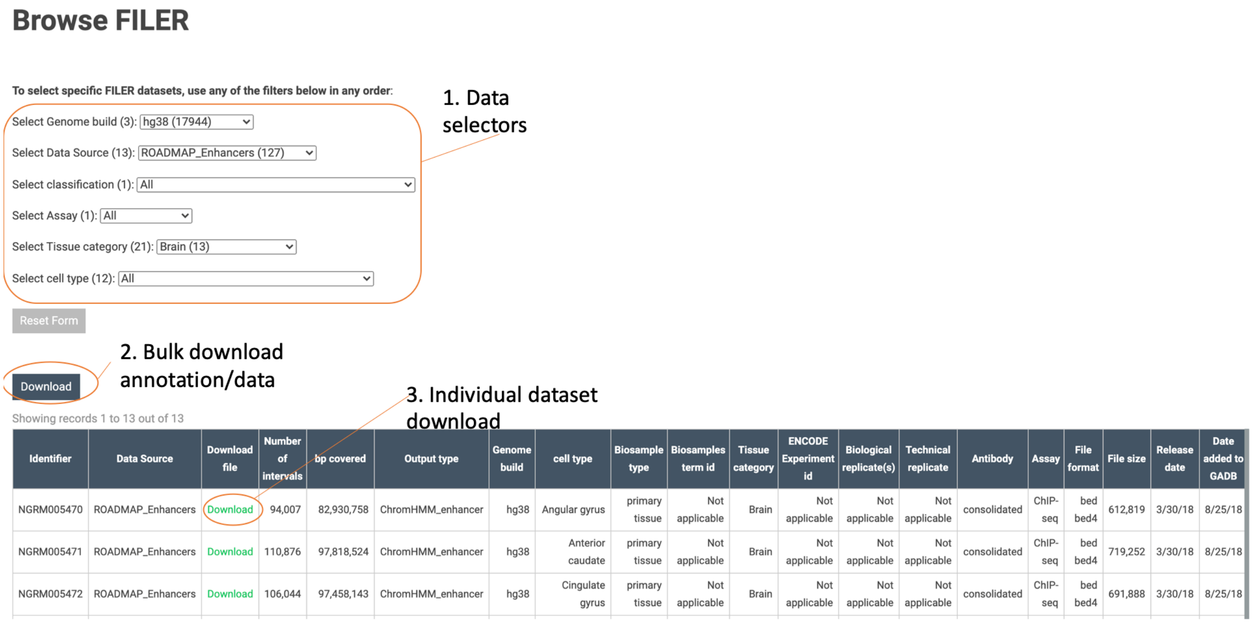


**Supplementary Figure S5**. Retrieving genomic and annotation datasets of interest using FILER.

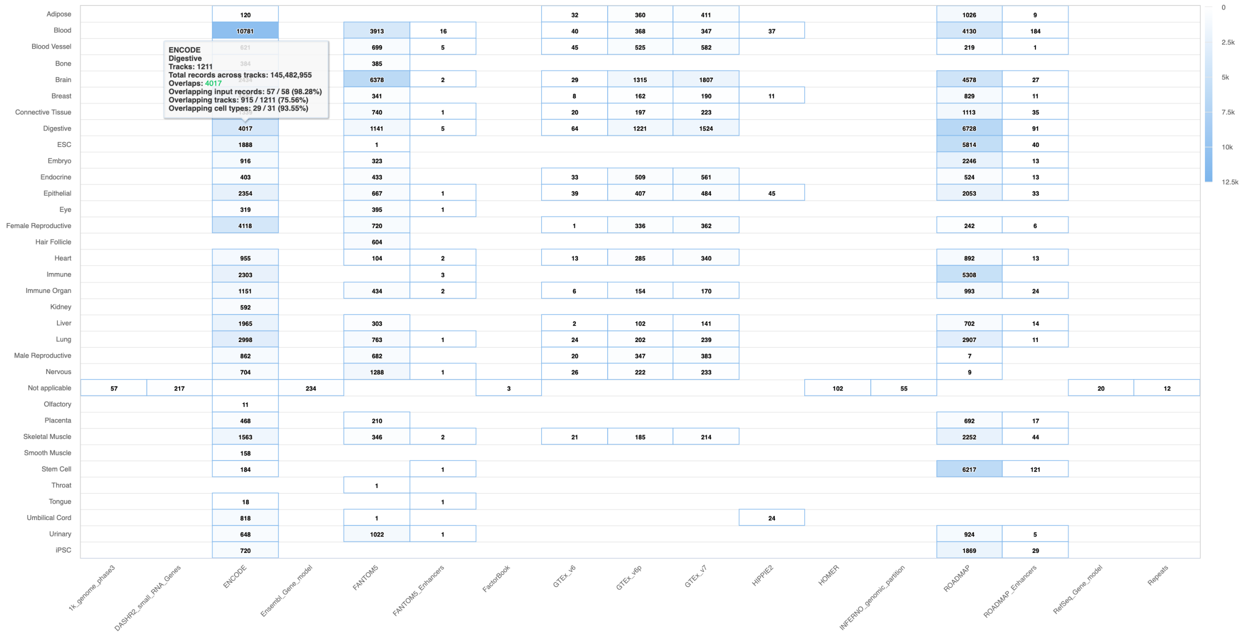


**Supplementary Figure S6**. Characterization of top genome-wide significant IBD variants. Shown is the summary of overlaps across tissue/cell type categories and data sources.

**Supplementary Table S1.** Summary of features for FILER and other similar tools

| **Features** | **FILER** | **Favor** (Li *et al.*, 2020) | **ENCODE Screen** (Davis *et al.*, 2018; Dunham *et al.*, 2012) |
|---|---|---|---|
| **Data integration** | | | |
| Cross-data source integration and harmonization of functional genomic data and metadata | >20 data sources | no | no (ENCODE only) |
| Wide biological context / biological categorization | >1,100 cell/tissue types; 36 tissue/cell type categories; 14 tissue and organ systems | no | 346 ENCODE cell/tissues types; 51 tissue categories |
| Unified data file formats | BED-based format | no | ENCODE only |
| Indexed and searchable functional genomics and annotation data | >50,000 datasets | No (variants only) | No (CRE[1] only) |
| Genome builds of data and annotation | GRCh38/hg38; GRCh37/hg19 | GRCh38/hg38 | GRCh38/hg38 |
| **Data analysis** | | | |
| Scalable interval-based genomic data search and annotation | Scalable genomic query engine | No (variants only) | No (CREs only) |
| Analysis and annotation of users' own data | Yes, SNPs or genomic regions | No (variants only) | No (overlap with CREs only) |
| Ability to use/query data with custom analysis pipelines | Web and command-line interface for data | No | No |
| **Data access** | | | |
| Web interface for genomic data search and query | Yes | No (variants only) | No (only CREs) |
| Scalable search / genomic data indexing | index-based genomic search | No | custom database (query CREs only) |
| Local (user-side) deployment | Yes (server, cloud, HPC[2] cluster) | No | No |
| Interactive filtering, search and download of genomic and annotation data | Yes (web interface) | No | No |

[1] CRE = candidate regulatory element

[2] HPC = high-performance computing cluster

**Supplementary Table S2.** Summary of integrated data sources
FILER_supplementary_Table_S2_data_source_summary_with_ref.xlsx

**Supplementary Table S3.** FILER data collections
FILER_supplementary_Table_S3_data_collections_summary.xlsx

**Supplementary Table S4.** FILER metadata schema description
FILER_supplementary_Table_S4_metadata_description.xlsx

**Supplementary Table S5.** Schema matching for integrated data sources
FILER_supplementary_Table_S5_schema_matching.xlsx

**Supplementary Table S6.** Data file formats/schemas for data tracks in the FILER database
FILER_supplementary_Table_S6_file_formats.xlsx

**Supplementary Table S7.** Summary of FILER data across biological annotation/data categories
FILER_supplementary_Table_S7_biological_data_category_summary.xlsx

**Supplementary Table S8.** FILER metadata table
FILER_supplementary_Table_S8_metadata.zip

**Supplementary Table S9**. Example of FILER annotated overlaps
FILER_supplementary_Table_S9_annotated_overlaps_example.xlsx

**Supplementary Note S1.** FILER data collection and preparation
FILER_supplementary_Note_S1_data_collection_and_preparation.docx
