## Supplementary Note S1. FILER data collection and preparation for "FILER: large-scale, harmonized FunctIonaL gEnomics Repository"

**FILER data collection, annotation and pre-processing**

This document provides detailed information on data collection, data pre-processing steps for each of data sources integrated into FILER.

**FILER metadata:**
Refer to “**FILER_supplementary_schema_matching.xlsx”** file for how and from where we obtained meta information for FILER

**Scripts**:

All scripts are accessible through FILER bitbucket repository:

[**http://bitbucket.org/wanglab-upenn/FILER**](http://bitbucket.org/wanglab-upenn/FILER)

**Pre-processing:**

All of the data provided in FILER are in BED format (other than 1000 genome VCF and genome sequence FASTA files). Refer to **“FILER_supplementary_file_formats.xlsx”** for different BED formats

BED files are 0-based sorted and indexed using GIGGLE and Tabix.

**Pre-processing scripts:**
Below scripts were used for every data source.

1) BED sort and bgzip (BED_sort-bgzip.sh)
2) Giggle indexing (GIGGLE_index.sh)
3) Tabix indexing (tabix_index.sh)

Other pre-processing steps that are specific for a particular data source are described in the documentation part below, along with the scripts used.

**ENCODE data collection, annotation, and pre-processing**

**Version**: Downloaded datasets released on or before 06/2020

**Data download**:

Access data from the experiment data matrix: <https://www.encodeproject.org/matrix/?type=Experiment&status=released>

Use the following filter terms:

Assay title: CHIP-seq, Dnase-seq, eCLIP, CAGE, FAIRE-seq, RNA-PET, ChIA-PET, RIP-seq, iCLIP, 5C, ATAC-seq, RNA-microarray [transcription profiling by array assay], CGH, RRBS, RIP-Chip, Switchgear, RAMPAGE, GM DNase-seq, microRNA counts

Genome assembly: hg19, hg38

Available file types: bigBed narrowPeak, bigBed broadPeak, bigBed bedRnaElements, bigBed tss_peak, bigBed idr_peak, bigBed bed12, bigBed bed3, gtf, gff gff3

After making the filter selection, use batch download (below instructions will be shown on ENCODE matrix page)

*Click the “Download” button download a “files.txt” file that contains a list of URLs to a file containing all the experimental metadata and links to download the file. The first line of the file has the URL or command line to download the metadata file.*

*The “files.txt” file can be copied to any server.
The following command using curl can be used to download all the files in the list:*

*xargs -L 1 curl -O -L < files.txt*

**Data pre-processing**:

1) Convert bigBed to BED (**bigBedToBed**)
2) Convert gff3 to BED (**GFF3toBed.sh**)
3) Convert gtf to BED (**GTFtobBed.sh**)

**GTEx data collection, annotation, and pre-processing**

**Version:** v6, v6p, v7, v8

**Data download**:

**GTEx v6:** significant snp-gene association
<https://storage.googleapis.com/gtex_analysis_v6/single_tissue_eqtl_data/GTEx_Analysis_V6_eQTLs.tar.gz>

**GTEx v6P:** significant snp-gene association
<https://storage.googleapis.com/gtex_analysis_v6p/single_tissue_eqtl_data/GTEx_Analysis_v6p_eQTL.tar>

**GTEx v6P:** all snp-gene association
<https://storage.googleapis.com/gtex_analysis_v6p/single_tissue_eqtl_data/GTEx_Analysis_v6p_all-associations.tar>

**GTEx v7**: significant snp-gene association
<https://storage.googleapis.com/gtex_analysis_v7/single_tissue_eqtl_data/GTEx_Analysis_v7_eQTL.tar.gz>
**GTEx v7**: all snp-gene association
<https://storage.googleapis.com/gtex_analysis_v7/single_tissue_eqtl_data/GTEx_Analysis_v7_eQTL_all_associations.tar.gz>

**GTEx v8 eQTL:** significant snp-gene association
<https://storage.googleapis.com/gtex_analysis_v8/single_tissue_qtl_data/GTEx_Analysis_v8_eQTL.tar>

**GTEx v8 eQTL:** all snp-gene association
Follow the instructions to download using google cloud
<https://console.cloud.google.com/storage/browser/gtex-resources/GTEx_Analysis_v8_eQTL_all_associations/>

**GTEx v8 sQTL:** significant snp-gene association
<https://storage.googleapis.com/gtex_analysis_v8/single_tissue_qtl_data/GTEx_Analysis_v8_sQTL.tar>

**GTEx v8 sQTL:** all snp-gene association
Follow the instructions to download using google cloud
<https://console.cloud.google.com/storage/browser/gtex-resources/GTEx_Analysis_v8_sQTL_all_associations/>

**Data pre-processing:**

**All snp-gene association –** Considering the file size we filtered variants with p-value < 0.05 script used for filtering - **“GTEx_filter_all_association.sh”**

Each version of GTEx has different formatting structure, we used GTEx v6p all snp-gene association as default format. Below scripts are used for creating BED files from GTEx data (conversion done after filtering variants for every all snp-gene association dataset)

1) GTEx_v6_eQTL_signif_bed_conversion.sh
2) GTEx_v6p_eQTL_signif_bed_conversion.sh
3) GTEx_v6p_eQTL_all_association_bed_conversion.sh
4) GTEx_v7_eQTL_signif_bed_conversion.sh
5) GTEx_v7_eQTL_all_association_bed_conversion.sh
6) GTEx_v8_eQTL_signif_bed_conversion.sh
7) GTEx_v8_eQTL_all_association_bed_conversion.sh
8) GTEx_v8_sQTL_signif_bed_conversion.sh
9) GTEx_v8_sQTL_all_association_bed_conversion.sh

After conversion to BED format, we rearranged the columns for consistency (to match v6p all snp-gene association) Script used for re-formatting - **“GTEx_BED_column_reformatting.sh”**

**ROADMAP data collection, annotation, and pre-processing**

**Version:** Release 9

**Data download:**

BroadPeak:
<https://egg2.wustl.edu/roadmap/data/byFileType/peaks/unconsolidated/broadPeak/>

GappedPeak:
<https://egg2.wustl.edu/roadmap/data/byFileType/peaks/unconsolidated/gappedPeak/>

NarrowPeak:
<https://egg2.wustl.edu/roadmap/data/byFileType/peaks/unconsolidated/narrowPeak/>

ChromHMM:
<https://egg2.wustl.edu/roadmap/data/byFileType/chromhmmSegmentations/ChmmModels/coreMarks/jointModel/final/>

i) all_hg38lift.mnemonics.bedFiles.tgz
ii) all.mnemonics.bedFiles.tgz

**Data pre-processing:**

1) ROADMAP enhancers – extracted enhancers from ChromHMM data – script used **“ROADMAP_enhancer_extraction.sh”**

**FANTOM5 data collection, annotation, and pre-processing**

**Version:** FANTOM5 Phase2.0

**Data download:**

Download the CAGS TSS bed files
http://fantom.gsc.riken.jp/5/datafiles/latest/basic/

README files are located inside every sub-folder

Enhancer download:
<http://slidebase.binf.ku.dk/human_enhancers/presets/serve/facet_expressed_enhancers.tgz>

**FactorBook**

**Version:** Release date: 16-March-2014

**Data download:**

<http://hgdownload.cse.ucsc.edu/goldenPath/hg19/database/factorbookMotifPos.txt.gz>

**Homer**

**Version:** 17/09/2017

**Data download:**

<http://homer.ucsd.edu/homer/data/motifs/>

**Human hg38 UCSC BigBed Track 170917:** <http://homer.ucsd.edu/homer/data/motifs/homer.KnownMotifs.hg38.170917.bigBed.track.txt>

**Human hg19 UCSC BigBed Track 170917:** <http://homer.ucsd.edu/homer/data/motifs/homer.KnownMotifs.hg19.170917.bigBed.track.txt>

**Data pre-processing:**

1) HOMER data has identical motif names, use script **“HOMER_modification_update_duplicate_motif_names.py**” to fix it

**TargetScan**

**Version:** 7p2 (March 2018)

**Data download:**

<http://www.targetscan.org/cgi-bin/targetscan/data_download.vert72.cgi>

**All predictions for representative transcripts**: Genome (hg19) locations of all targets, partitioned into files by conservation of miRNA family and site

**Default predictions (conserved sites of conserved miRNA families):** Genome (hg19) locations of human predicted (conserved) targets of conserved miRNA families

**Reference Genome**

**hg19**: <http://hgdownload.soe.ucsc.edu/goldenPath/hg19/chromosomes/>
Feb. 2009 assembly of the human genome

**hg38**: <http://hgdownload.soe.ucsc.edu/goldenPath/hg38/chromosomes/>
Dec. 2013 assembly of the human genome

**Gene model**

**Ensembl**

**hg19:** <ftp://ftp.ensembl.org/pub/release-75/gtf/homo_sapiens>
release 75

**hg38**: <ftp://ftp.ensembl.org/pub/release-89/gtf/homo_sapiens>
release 89

**RefSeq**
Used UCSC table browser to download NCBI RefSeq data

**hg19:** <http://genome.ucsc.edu/cgi-bin/hgTables>
genome: Human, assembly: Feb. 2009 (GRCh37/hg19), group: Genes and Gene Predictions, Track: NCBI RefSeq

**hg38**: <http://genome.ucsc.edu/cgi-bin/hgTables>
genome: Human, assembly: Dec. 2013 (GRCh38/hg38), group: Genes and Gene Predictions, Track: NCBI RefSeq

**Repeats**

**hg19:** <http://genome.ucsc.edu/cgi-bin/hgTables>
genome: Human, assembly: Feb. 2009 (GRCh37/hg19), group: Repeats, Track: RepeatMasker

**hg38:** <http://genome.ucsc.edu/cgi-bin/hgTables>
genome: Human, assembly: Dec. 2013 (GRCh38/hg38), group: Repeats, Track: RepeatMasker

**DASHR2 data**

**Version:** DASHR2 (August-2018)

**Data Download:**

**Small RNA called peaks data:**

<http://dashr2.lisanwanglab.org/download.php>

Select DASHR data collection: Download sncRNA tables (annoted and unnotated from Small RNA loci table row)

<http://dashr2.lisanwanglab.org/table2csv.php?dataSourceDownload=ENCODE_GEO_hg19&table=peaks>

<http://dashr2.lisanwanglab.org/table2csv.php?dataSourceDownload=DASHR2_GEO_hg38&table=peaks>

<http://dashr2.lisanwanglab.org/table2csv.php?dataSourceDownload=ENCODE_GEO_hg19&table=peaks>

<http://dashr2.lisanwanglab.org/table2csv.php?dataSourceDownload=ENCODE_GEO_hg38&table=peaks>

<http://dashr2.lisanwanglab.org/table2csv.php?dataSourceDownload=ENCODE_dataportal_hg19&table=peaks>

<http://dashr2.lisanwanglab.org/table2csv.php?dataSourceDownload=ENCODE_dataportal_hg38&table=peaks>

<http://dashr2.lisanwanglab.org/table2csv.php?dataSourceDownload=DASHR1_GEO_hg19&table=peaks>

<http://dashr2.lisanwanglab.org/table2csv.php?dataSourceDownload=DASHR1_GEO_hg38&table=peaks>

**DASHR Annotation:**

<http://dashr2.lisanwanglab.org/downloads/dashr.v2.sncRNA.annotation.hg19.gff>
<http://dashr2.lisanwanglab.org/downloads/dashr.v2.sncRNA.annotation.hg38.gff>
<http://dashr2.lisanwanglab.org/downloads/dashr.v2.annotation.hg19.gff>
<http://dashr2.lisanwanglab.org/downloads/dashr.v2.annotation.hg38.gff>

**Data pre-processing:**

Used script **“DASHR2_split_by_tissue.sh”** to split the peak files by tissues.

**1000Genome**

**Version:** Phase3

**Data Download:**

Download the vcf files from the following FTP links

Hg19: <ftp://ftp.1000genomes.ebi.ac.uk/vol1/ftp/release/20130502/>

Hg38: <ftp://ftp.1000genomes.ebi.ac.uk/vol1/ftp/release/20130502/supporting/GRCh38_positions/>

**Data pre-processing:**

VCF files for each chromosome were split by variant type (SNP, INDEL, biallelic, multiallelic), super and sub population.

Scripts used for splitting the VCF files are in the package **“1kgenome_pre-processing_scripts.tar.gz”**

**Hg38 liftover**

Liftover was performed on all of the data sources for which no hg38 data was available.

Hg19 genome coordinates are lifted using UCSC liftOver utility **hg19ToHg38.over.chain.gz**

Hg19 coordinates are retained in the lifted-over files (column 4, 5 and 6 are hg19 coordinates in the lifted-over files)

**Download link:** <http://hgdownload.cse.ucsc.edu/goldenPath/hg19/liftOver/hg19ToHg38.over.chain.gz>

<http://hgdownload.cse.ucsc.edu/admin/exe/linux.x86_64/liftOver>
